# Targeted immuno-antiretroviral HIV therapeutic approach to provide dual protection and boosts cellular immunity: A proof-of-concept study

**DOI:** 10.1101/2020.04.20.050849

**Authors:** Subhra Mandal, Shawnalyn W. Sunagawa, Pavan Kumar Prathipati, Michael Belshan, Annemarie Shibata, Christopher J. Destache

## Abstract

Human immunodeficiency virus (HIV)-infected active and latent CCR5 expressing long-lived T-cells are the primary barrier to HIV/AIDS eradication. Broadly neutralizing antibodies and latency-reversing agents are the two most promising strategies emerging to achieve ‘functional cure’ against HIV infection. Antiretrovirals (ARVs) have shown to suppress plasma viral loads to non-detectable levels and above strategies have demonstrated a ‘functional cure’ against HIV infection is achievable. Both the above strategies are effective at inducing direct or immune-mediated cell death of latent HIV+ T-cells but have shown respective limitations. In this study, we designed a novel targeted ARVs-loaded nanoformulation that combines the CCR5 monoclonal antibody and antiretroviral drugs (ARV) as a dual protection strategy to promote HIV ‘functional cure’. The modified CCR5 monoclonal antibody (xfR5 mAb) surface-coated dolutegravir (DTG) and tenofovir alafenamide (TAF) loaded nanoformulation (xfR5-D+T NPs) were uniformly sized <250 nm, with 6.5 times enhanced antigen-binding affinity compared to naïve xfR5 mAb, and provided prolonged DTG and TAF intracellular retention (t_1/2_). The multivalent and sustained drug release properties of xfR5-D+T NPs enhance the protection efficiency against HIV by approximately 12, 3, and 5 times compared to naïve xfR5 mAb, D+T NP alone, and xfR5 NPs, respectively. Further, the nanoformulation demonstrated high binding-affinity to CCR5 expressing CD4+ cells, monocytes, and other HIV prone/latent T-cells by 25, 2, and 2 times, respectively. Further, the xfR5-D+T NPs during short-term pre-exposure prophylaxis induced a protective immunophenotype, i.e., boosted T-helper (T_h_), temporary memory (TM), and effector (E) sub-population. Moreover, treatment with xfR5-D+T NPs to HIV-infected T-cells induced a defensive/activated immunophenotype i.e., boosted naïve, T_h_, boosted central memory, TM, EM, E, and activated cytotoxic T-cells population. Therefore, this dual-action targeted mAb-ARV loaded nanoformulation could potentially become a multifactorial/multilayered solution to achieve a “functional cure.”

## Introduction

Currently, antiretroviral therapy (ART) is the prime treatment strategy for human immunodeficiency virus (HIV) infection. ART improves HIV patient’s life-expectancy by effectively controlling plasma viral load (pVL), but is unable to eradicate the virus, thus patients have to commit to continuing life-long ART. Additionally, ART withdrawal could reactivate latent virus. Therefore, alternative ways are under investigation to search for potential candidates to ‘functionally-cure’ HIV infection (1). One of the essential targets of HIV research is the C-C motif chemokine receptor 5 (CCR5), a co-receptor that is predominant expressed on CD4+ T-cells, latently HIV-1 infected cells, dendritic cells (DC) and macrophages, and is responsible for HIV-1 entrance into the targeted cells (2).

Maraviroc, a CCR5 antagonist that blocks HIV entry and infection by docking on the CCR5 receptor on CCR5+ cells, is the first approved CCR5-antagonist drug for HIV-1 treatment (3). The other promising approach is genome editing that disrupts CCR5 alleles in CD4+ T-cells due to the infusion of an engineered zinc finger nuclease (ZFN) (4). However, the only gene-therapy approach to date that had conferred HIV cure is the use of CCR5 delta32 natural mutant genotype. It resists HIV-1 entry in three patients, i.e., the “Berlin Patient”, “London Patient”, and the “Düsseldorf patient” upon transplantation of stem cells from a CCR5 delta32 genotype donor (5).

Even though the sustained ART-free remission is a well-accepted and effective strategy to control HIV infection, the main focus of HIV research is to develop strategies that prolong protection and target latent HIV+ cells. Recent studies have demonstrated designing a long-acting (LA) ARV delivery system would boost the HIV prevention and treatment strategy (6–9). The LA ARV delivery system has shown to enhance drug-solubility, stability, biodistribution, pharmacokinetics, efficiency, and concurrent drug safety, due to reduced ARV associated side-effects, such as mitochondrial toxicity, renal abnormality, and reduced bone mineral density (10–12). Even though LA ARV nanoparticles (NPs) are promising to be a successful injectable LA antiretroviral candidate, these systems still need to show a good tolerability profile. In the LATTE-2 trial, the cabotegravir plus rilpivirine LA injection group has reported grade 3-4 adverse events higher compared to the oral comparative treatment group (13). The ECLAIR prevention study of LA ARVs (e.g., cabotegravir LA plus rilpivirine LA injection group) (14, 15) revealed a significantly prolonged sub-therapeutic tail of residual drugs which places patients at high-risk to contracting HIV and the possibility of developing resistance (16). The emergence of ARV resistance would limit future treatment options in those patients treated with LA ARV NPs. However, ‘targeting’ HIV prone cells is another strategy that is still under investigation.

The broadly neutralizing antibodies (bNAbs) against HIV have emerged as a promising strategy to protect or to treat circulating HIV (17). However, bNAbs for HIV is still a naïve field, as the search is on-going to find Abs that have good neutralization potency (18). Eliminating the circulating HIV is not sufficient to achieve “functional cure” against HIV, due to the presence of latently HIV-infected cells. Furthermore, the reported bNAbs still need optimization in terms of their effector function and plasma half-life (19–21). Therefore, the bNAbs strategy reveals that antibody-based HIV therapy has two limiting factors, i.e., plasma concentration and neutralization potency. Recently, studies have already confirmed the existence of resistant HIV-1 strains against bNAbs (22, 23), challenging the bNAbs clinical potency.

Furthermore, combination ARV (cARV) therapy could suppress plasma viral load but has been unsuccessful in immune restoration. Recovery from the viral infection needs reestablishment of strong immunity. Therefore, research against HIV-1, faces challenges related to immune reconstitution failure. Studies have revealed that the HIV infection induce terminal differentiation of effector (E) CD8+ T cells to the memory phenotypes that causes progressive E population reduction, immune exhaustion, and promote activation-induced cell death (24). HIV-infection strongly compromises the immune system and immune impairment results in its inability to respond to HIV and other pathogens; consequently, rapid progression to acquired immunodeficiency syndrome (AIDS).

However, ideally, the immune reconstitution could be promoted by stimulating both humoral immune responses and cellular immune responses to prevent and control HIV infection, respectively. The adaptive immune system plays the most critical role in protection against HIV-1 infection. Studies have shown that promoting T-cell-based immunity, specifically cytotoxic T lymphocyte (CTL) stimulation (25, 26), would promote effective production of neutralizing antibodies, and would affect synergistically to protection against active and latent HIV-1 infection.

Our hypothesis is a combination of HIV therapeutic strategies, i.e., ART and antibody-mediated HIV-entry blockage, would be an effective strategy to suppress HIV viremia, and promote protective immune-response. The cumulative effect could potentially induce a “functional cure.” In this study, we propose an innovative approach to combine ART and CCR5 mAb in a novel single nano-module to achieve dual protection against HIV infection. To achieve this, a novel LA ARV loaded and anti-CCR5 mAbs surface decorated nanoparticles (NPs) have been formulated (Figure 1). The first level of protection comes from CCR5 mAb (decorated on the ARV NP surface), that blocks HIV-1 entry in the CD4+ cells surface (i.e., CCR5+ CD4+ cells) similar to the CCR5 antagonist. Wherein, internalization of CCR5/ CCR5 mAb-ARVs NP complex causes release and maintaining intracellular ARV, providing the second level of protection again HIV infection (Figure 1). Our study shows the novel CCR5 mAb decorated ARV nanoformulation act as a single delivery-system with dual-layers of defense that could prevent primary HIV-infection as well as potentially induce a protective immunophenotype, to target and suppress HIV latency. Therefore, a dual protection mechanism will prevent new HIV infection of the naive and latent CCR5+ cell population, potentially achieving a “functional cure.”

**Figure 1.**
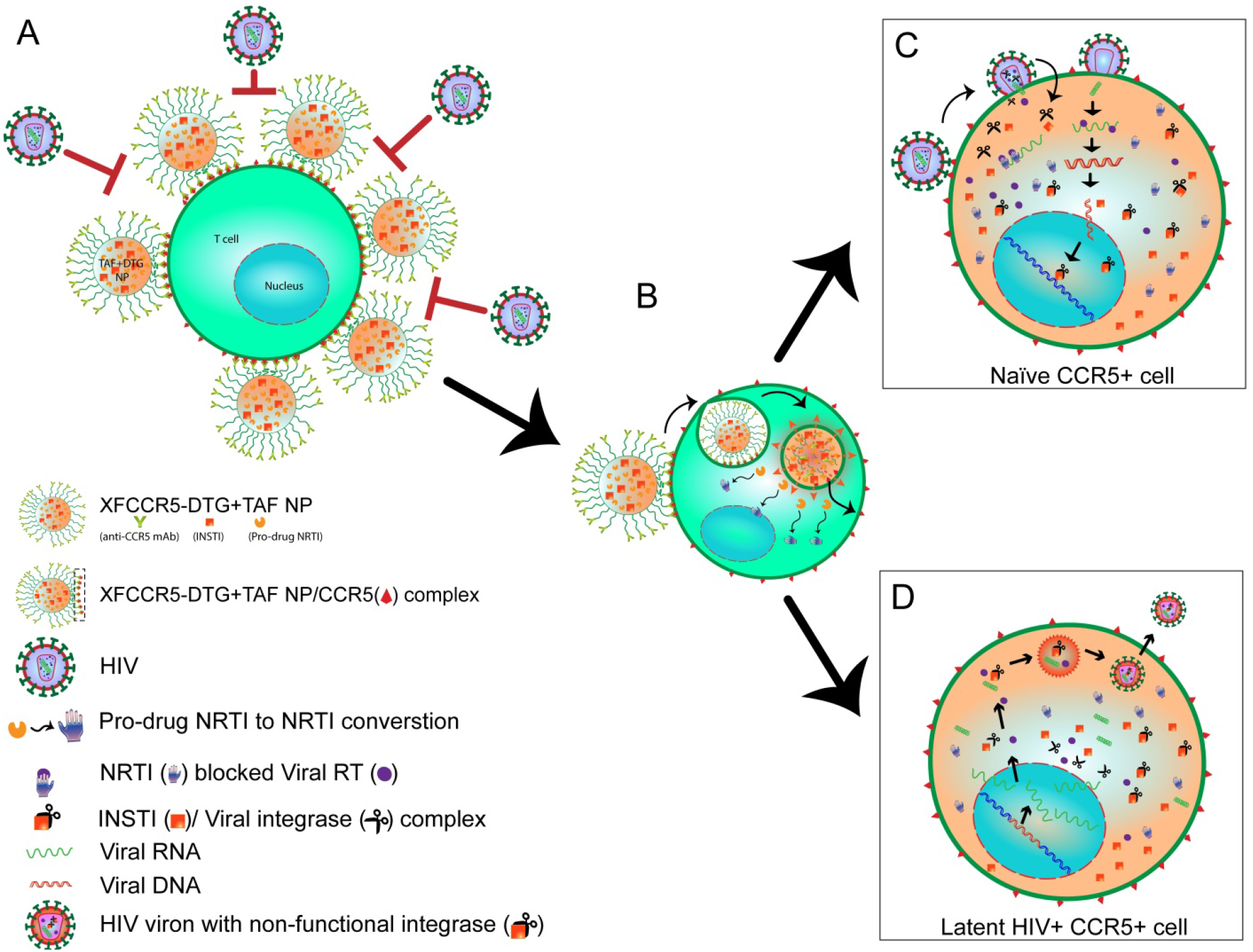
Schematic diagram explaining the dual-action strategy of targeted cARV-nanoformulation. A) First level of protection: xfR5 D+T NP bound on the T-cell surface via CCR5 receptor (xfR5-D+T NP/CCR5 complex) blocks HIV interaction with CCR5 on T-cell surface by two approaches, i.e., first, by blocking HIV from CCR5 binding and the second, providing steric hindrance to HIV virions due to nanoformulation crowding on T-cell surface (red block arrows) (65). B) The xfR5 D+T NP/CCR5 complex triggers the CCR5 trafficking pathway due to anti-CCR5 mAb binding with the CCR5+ T-cells (66), leading to internalization of xfR5 D+T NP. Within the endosome, the low pH induces degradation of polymeric NPs, leading to the release of DTG (INSTI)and TAF (NRTI) into the cytoplasm. The second level of protection: the presence of an intercellular pool of TFV di-phosphates and DTG to prevent (C) naive cell HIV infection. (D) The latently HIV+ cells, free intracellular high-affinity DTG (INSTI), will bind with fresh integrase enzymes produced during the reactivation stage, resulting in a non-functional INSTI/integrase complex. Therefore fresh virons thus produced will be with non-functional INSTIs. Therefore, fresh viron will not be able to integrate viral DNA in the host genome.

## Results

### CCR5 targeted cARV loaded NPs characterization

The CCR5 targeted cARV loaded NPs were formulated by using standardized methodology with some modification (9, 10). To conjugate targeting molecule on ARV loaded NP’s surface, NHS functionalized NP loaded with DTG+TAF were formulated (Figure 1). The modified water-in-oil-in-water (w-o-w) interfacial polymer deposition method leads to well-defined N–hydroxysuccinimide (NHS) functionalized DTG+TAF NP (NHS-D+T NP) formulation. The DLS analysis demonstrated D+T NPs obtained averaged 198.7 ± 10 nm with uniform size-distribution pattern (PDI: 0.153 ± 0.008) (Table 1). Moreover, the modified CCR5 mAb (xfR5 mAb) conjugated D+T NP (xfR5-D+T NP) demonstrated a slight increase in size to 212.6 ± 20.7 nm. The shape and surface properties evaluation by scanning electron microscopy reflected that NHS-D+T NPs obtained were uniform, and smooth-surfaced spherical particles (Figure 2A) and xfR5 mAbs surface conjugation does not cause any change in the particle’s morphology (Figure 2B). The xfR5 mAb binding reduced the surface charge density of D+T NPs (−28.15 ± 1.9), to −17.77 ± 1.9 for xfR5-D+T NPs (Table 1).

**Figure 2.**
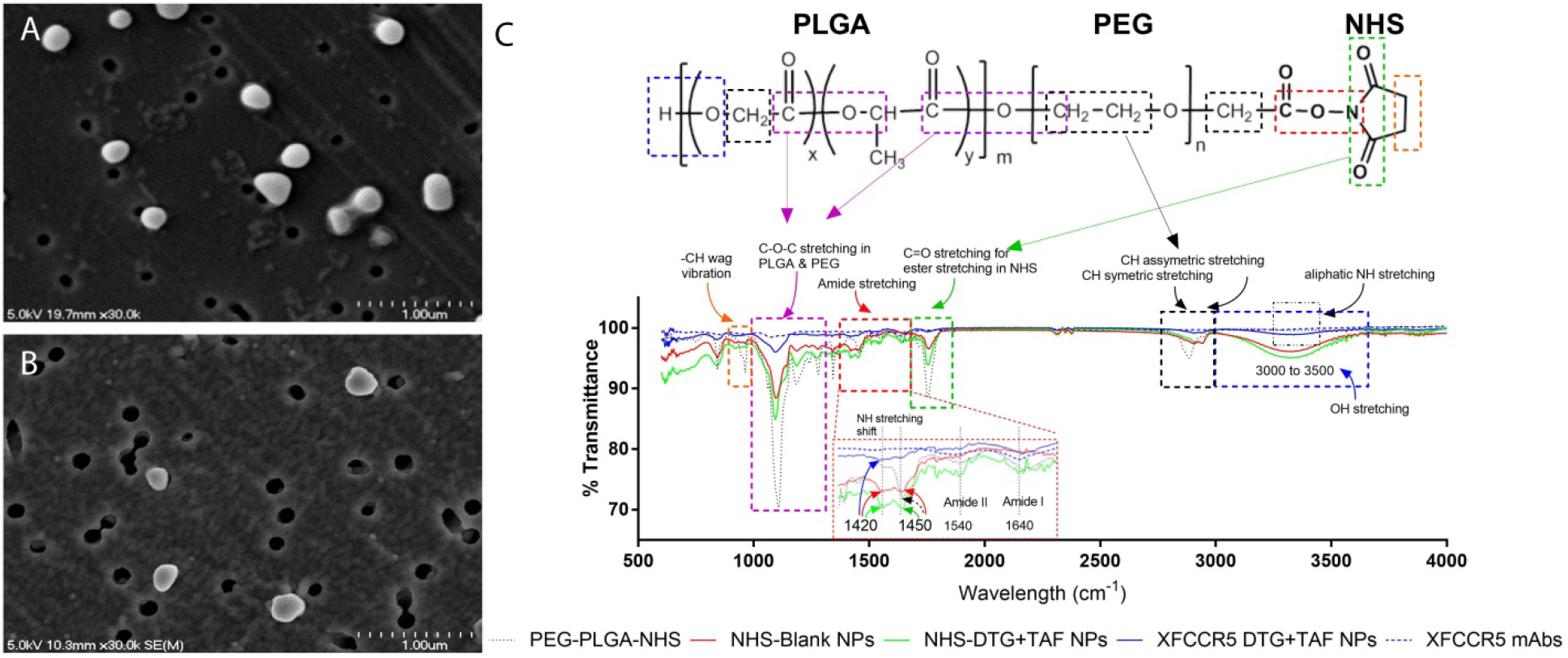
Representative scanning electron microscopic image of NHS-D+T NP (A) and xfR5 D+T NP (B). C) Representative FT-IR spectrum obtained from PEG-PLGA-NHS polymer, NHS functionalized NPs (NHS-Blank NPs), NHS functionalized D+T NPs (NHS-D+T NPs), xfR5 D+T NPs, and xfR5 mAbs. The significant functional group bands are presented in various color boxes (Orange: –CH wag and rocking vibration band (840-900 cm^-1^); Purple: C-O-C stretching bands (1095-1198 cm^-1^); Red: amide bond; Green: C=O stretching band (1695-1818 cm^-1^); Red: −CON-stretching based on reported FT-IR spectral studies (67, 68). Inset graph enlargement image (C) shows NH-band shifting from 1450 cm^-1^ to 1420 cm^-1^ indicating COO-NC bond conversion to amide bond (–CONH–); and presence of Amide I and II at 1540 cm^-1^and 1640 cm^-1^ respectively.

**Table 1.** Physicochemical characteristics of D+T NPs and xfR5-D+T NPs.

| Type | Size (nm) | Surface charge (mV) | Polydispersity Index (PDI) | %Drug entrapment efficiency (EE) | xfR5 mAb bound per mg D+T NP |
| --- | --- | --- | --- | --- | --- |
| NHS-D+T NP | $198.7 \pm 10$ | $-28.15 \pm 1.9$ | $0.153 \pm 0.008$ | DTG: $58.8 \pm 10$<br>TAF: $60.5 \pm 10.3$ | N/A |
| xfR5-D+T NP | $212.6 \pm 20.7$ | $-17.77 \pm 1.9$ | $0.203 \pm .0175$ | N/A | $3.7 \pm 0.52 \mu\text{g}$ |

*Data represented as mean ± SEM, obtained from three independent batches of NHS-D+T NPs or xfR5-D+T NPs (n=3); N/A= not applicable*

This reduction of the surface charge could be attributed to several factors such as charge masking by the ions from the PBS buffer or to xfR5 mAb surface decoration or both. The polymeric NPs encapsulation resulted in enhanced drug encapsulation efficiency (% EE) of 58.5 ± 5.2 % and 55.7 ± 3.9 % for DTG and TAF, respectively (Table 1). Therefore, DTG and TAF were loaded at a 1:1 ratio in the xfR5-D+T NPs.

The xfR5 mAb were conjugation on NHS-D+T NP by carbodiimide-crosslinking mechanism (27). The xfR5 mAb binding on xfR5-D+T NPs quantified by BCA assay, estimated ~ 3.7 ± 0.52 μg of xfR5 mAb per mg xfR5-D+T NPs (Table 1). The xfR5 mAb binding was further confirmed by FT-IR analysis, as shown in Figure 1C. The typical NHS ester bands at 1695-1818 cm^-1^ (aromatic C=O stretching, green dashed-line box) and 967 cm^-1^ (–CH wagging band, orange dashed-line box) of PEG-PLGA-NHS polymer, on NHS-Blank NPs and NHS-D+T NPs, confirmed the presence of free NHS functional group on the NP surface. The amide bond obtained when the primary amine group of the antibody by replacing the −NHS group of NHS-PEG on NPs surface (28). Thus, the disappearance of these −NHS band in xfR5-D+T NP proved the replacement of NHS ester group. Additionally, the shift of primary aliphatic amine NH stretching (3350 cm^-1^ and 1420 cm^-1^), and −CH stretching (2947 cm^-1^) (29), along with primary amide I and II band presence (at 1640 and 1540 cm^-1^ respectively) confirmed the covalent amide (-CONH-) bond formation. Whereas, the presence of other significant bands C-O-C stretching (1000-1300 cm^-1^) and broad O–H stretching band (3000-3500 cm^-1^) confers xfR5-D+T NPs surface has open PLGA & PEG surfaces along with covalently bound xfR5 mAb. Therefore, xfR5 mAb are not densely packed on xfR5-D+T NP and have enough free space to avoid steric hindrance during targeted CCR5 receptor binding.

### Binding Affinity

The multimeric xfR5 mAb on xfR5-D+T NP enhance the binding affinity compared to xfR5 mAb alone. Thus, the antibody binding affinity was estimated by flow cytometric analysis, and the data are expressed as equilibrium dissociation constant, K_d_. The binding affinity (K_d_) of xfR5-D+T NP compared to naïve xfR5 mAb and wild-type CCR5 mAb on TZM-bl cells (CCR5+ CD4+ cell line) (Table 2), demonstrates xfR5 mAb displayed approximately 10-fold higher affinity (0.251 ± 0.15 nM) compared to wild-type CCR5 mAb (2.023 ± 0.655 nM). Simultaneous multiple xfR5 mAb within the xfR5-D+T NP (5.7 ± 2.7 nM) binding further improved the sensitivity by approximately 6.6 times (Supplementary Figure 1 and Table 2).

**Table 2.** Comparative binding affinity (K_d_) of xfR5-D+T NPs vs. xfR5 mAb with TZM-bl cells (CD4+ CCR5+ cell line), primary CD4+ T-cells (T_h_ cells), and with other HIV-1 prone and CCR5 receptor-expressing immune cells (primary CD8+, CD68+, and CD2+ T-cells).

| Type | Binding Affinity ($K_d$, nM) | | | | |
| --- | --- | --- | --- | --- | --- |
|  | TZM-bl cells | CD4+ T-cells | CD8+ T-cells | CD68+ T-cells | CD2+ T-cells |
| xFR5-D+T NPs | $0.038 \pm 0.020$ | $4.31 \pm 1.47$ | $0.99 \pm 0.38$ | $0.11 \pm 0.066$ | $1.5 \pm 1.1$ |
| xFR5 mAbs | $0.251 \pm 0.15$ | $20.87 \pm 10.65$ | $25.02 \pm 14.3$ | $0.21 \pm 0.097$ | $2.6 \pm 2.5$ |
| Wild-type anti-CCR5 mAb | $2.023 \pm 0.655$ | ND | ND | ND | ND |

*Data represented as mean ± SEM obtained from three healthy independent donors; ND= not determined.*

Similarly, xfR5-D+T NPs (4.31 ± 1.47 nM) resulted in enhanced binding with the primary CD4+ T-cells (prime HIV-1 infecting CCR5 expressing T-cell type), i.e., approximately 5 times lower K_d_ value compared to xfR5 mAb alone (20.87 ± 10.65 nM).

The binding affinity of xfR5-D+T NPs with other CCR5+ immune cell types and HIV-prone cells such as cytotoxic T-cells (CTLs, CD8+), dendritic T-cells (CD2+) and monocytes (CD68+), were evaluated in similar a method (gating strategy, supplementary Figure 2). The comparative binding affinity study demonstrated xfR5-D+T NPs had an enhanced binding affinity with memory CD8+ T-cells, approximately 25 times higher affinity compared to naïve xfR5 mAb (Table 2). Whereas xfR5-D+T NP binding with CD2+ T-cells and CD68+ T-cells illustrated slightly enhanced but non-significant difference in binding affinity compared to naïve xfR5 mAb.

### Intracellular drug kinetics

The sustained release property of xfR5-D+T NP compared to D+T in solution was evaluated by evaluating the intracellular uptake/release and retention kinetics based on the non-compartmental analysis using Phoenix WinNonlin 8.1 software (Table 3).

**Table 3.** Comparative intracellular kinetics study of DTG, TAF, active drug (TFV) and its metabolite (TFV-dp), upon xfR5-D+T NPs or D+T solution treatment.

| Parameter | Units | NP |  |  |  | Solution |  |  |  |
| --- | --- | --- | --- | --- | --- | --- | --- | --- | --- |
|  |  | TAF | TFV | DTG | TFV-dp | TAF | TFV | DTG | TFV-dp |
| $C_{max}$ | (pmole/<br>10<sup>6</sup><br>cells) | 18.1<br>±3.3 | 1250.2<br>±269.1 | 37027.8<br>±5401.0 | 29.7<br>±13.1 | 1.4±<br>0.7 | 1428.4<br>±580.4 | 334.0±<br>197.4 | 12.9±<br>4.1 |
| $AUC_{all}$ | h*(pmole/<br>10<sup>6</sup><br>cells) | 825.<br>4±11<br>9.5 | 52384.<br>8±4613<br>.2 | 224049<br>0.8±<br>240106.<br>3 | 827.<br>5±12<br>5.5 | 11.4<br>±4.4 | 47465.<br>0±9641<br>.7 | 8805.0<br>±1934.<br>1 | 585.9<br>±94.6 |
| $t_{1/2}$ | h | 75.5 | 21.6 | 79.2 | 32.3 | 6.5 | 22.3 | 18.0 | 25.4 |

$C_{max}$ = maximum concentration; $AUC_{all}$ = area under the curve; $t_{1/2}$ = half-life of the drug dose; Data presented as mean ± SEM of three independent experiments (n=3), with each experiment with three repeats/variable.

The D+T NP compared to the D+T solution demonstrated that the concentration maximum (C_max_) and the area under the concentration-time curve (AUC_all_) of TAF from xfR5-D+T NP were 12.9 and 72.4 times higher than TAF in solution. In contrast, DTG demonstrated comparatively higher C_max_ and AUC_all_, (110.9 and 254.5 times higher), respectively. Whereas, these parameters showed a non-significant difference for intracellular TFV and TFV-dp, between xfR5-D+T NP and D+T solution treatment. The rationale behind this non-significant difference in the active-drug, TFV and TFV-dp could be due to its dependence on cellular enzyme kinetics. The conversion of TAF >TFV >TFV-dp intracellularly, is restricted due to acquired steady state. Thus, in the absence of intracellular drug utilization, the cellular TFV and TFV-dp steady-state concentration are maintained in both case of xfR5-D+T NP and D+T solution treatments. From the Cmax and AUCall data of TAF and DTG, it is evident that the nanoformulation enhanced cellular uptake of ARV compared to the same drugs in solution. In terms of retention, NP demonstrated 11.6 and 4.4 times higher TAF and DTG elimination half-life (t_1/2_), than naïve drugs in solution, which is indicative of improved retention kinetics.

### Cytotoxicity and HIV Protection study

The nanoencapsulation of ARV drugs has known to reduce the toxicity of naïve drugs (9, 10, 30, 31). A comparative cytotoxicity study performed on TZM-bl cells and primary peripheral blood mononuclear cells (PBMCs) (Table 4).

**Table 4.** Comparative cytotoxicity (CC_50_), protection (IC_50_), and selectivity index (SI, CC_50_/ IC_50_) study of xfR5-D+T NPs vs. xfR5 mAb on TZM-bl cells and primary CD4+ T-cells.

| Cell Type | Treatment Type | $CC_{50}$ (nM) | $IC_{50}$ (nM) | SI |
| --- | --- | --- | --- | --- |
| TZM-bl | xfR5-D+T NPs | 2910 ± 134.9 | 0.0352 ± 0.0086 | 82670 |
|  | xfR5 NPs | 998.7 ± 122.2 | 0.05471 ± 0.0103 | 18254 |
|  | D+T NPs | 3095 ± 102.3 | 0.195 ± 0.16 | 15872 |
|  | xfR5 mAb | 1464 ± 35.5 | 18.53 ± 2.85 | 79 |
| PBMCs | XF CCR5 D+T NPs | 690.6 ± 114.7 | 9.77 ± 2.03 | 71 |
|  | XF CCR5 NPs | 1428 ± 83.5 | 53.34 ± 2.34 | 26 |
|  | D+T NPs | 1009 ± 88.7 | 33.95 ± 1.91 | 30 |
|  | XF CCR5 mAb | 816.7 ± 93 | 115.9 ± 1.61 | 7 |

*Each data represented as mean ± SEM (n=3, healthy independent donors); CC<sub>50</sub>=50% cytotoxic concentration; IC<sub>50</sub>= half-maximal inhibitory concentration*

The cytotoxicity on TZM-bl cells, demonstrated xfR5-D+T NP have no toxicity compare to xfR5 mAb. The study revealed all the tested nanoformulations (xfR5-D+T NP, xfR5 NP, and D+T NP) were safe for cellular application. Moreover, xfR5-D+T NP demonstrated 2-times higher 50% cytotoxic concentration (CC_50_) value (2910 ± 134.9 nM) compared to xfR5 mAb (1464 ± 35.5 nM). However, in primary PBMCs, i.e., xfR5-D+T NP (690.6 ± 114.7 nM) and xfR5 mAb (816.7 ± 93 nM) treatment demonstrated non-significant CC_50_ differences (Table 4). The *in vitro* results on TZM-bl cells and PBMCs suggest that nano-encapsulation reduces the toxic effect of DTG and TAF as well as promotes cell viability.

The dual protection by xfR5-D+T NP due to multimeric xfR5 mAb blocking and prolonged ARV release is expected to improve the half-maximal inhibitory concentration (IC_50_), compared to xfR5 mAb and D+T NPs alone. The HIV protection efficacy study in TZM-bl cells demonstrated xfR5-D+T NP (0.0352 ± 0.0086 nM) improved IC_50_ by 528 and 5.5 times compared to D+T NPs (0.195 ± 0.1 nM) and xfR5 mAb (18.53 ± 2.85 nM), respectively (Table 4). Similarly, in primary PBMCs, where only a fraction of cell expresses CCR5 surface receptor, xfR5-D+T NP treatment improved the protection against HIV-infection by 12, 5.5 and 3.4 times compared to naïve xfR5 mAb, xfR5 NP, and D+T NP, respectively (Table 4). Further, the xfR5 NP (with ARVs) reduced IC_50_ by 338 times (TZM-bl: 0.05471 ± 0.0103 nM) and 2-times (PBMCs: 53.34 ± 2.34 nM) compared to naïve xfR5 mAb. This result supports the hypothesis that multimeric mAb coated nanoformulation induces multi-valent interaction receptor/ligand interaction and thus enhances protection efficacy compared to xfR5 mAb alone (Supplementary Figure 1) (32, 33).

Therefore, the selectivity index (SI) evaluation significantly reflected the potential of xfR5-D+T NPs (Table 4). In TZM-bl cells (all cells uniformly CCR5+ CD4+ cells), the xfR5-D+T NP improves SI value by 5.5 and 4.5 times higher than individualized treatment by xfR5 NP and D+T NPs. However, in primary PBMCs, xfR5-D+T NP demonstrated 2.7, 2.4, and 11 times higher SI value compared to xfR5 NP, D+T NP, and naïve xfR5 mAb, respectively. These results showed xfR5 decorated D+T NP, significantly improved the therapeutic index of both individual therapeutic approaches, i.e., xfR5 mAb and D+T NPs.

### Immunophenotype during *in vitro* short-term PrEP and HIV-1 treatment study

Immunophenotype performed on PBMCs isolated from healthy donors to evaluate the immunological potential of targeted ARV-loaded nanoformulation. The immune-differentiation pattern of T-cells during PrEP and HIV-1 treatment was evaluated by flow cytometry after treating uninfected PBMCs with xfR5-D+T NP or xfR5 mAb, along with respective controls. Supplementary figure 3 illustrates the gating strategy for the immunophenotypic study. Based on the expression level of CCR7, CD27 and CD45RO receptors on T-cells, five distinct T-cell sub-populations were determined, i.e., naïve cells (CCR7+ CD27+ CD45RO-), central memory cells (CM, CCR7+ CD27+ CD45RO+), transition memory cells (TM, CCR7-CD27+ CD45RO+), effector memory cells (EM, CCR7-CD27-CD45RO+), and effector cells (CCR7-CD27-CD45RO-). The immunophenotype of the T-cell sub-populations were determined over different time-points and conditions, day before PHA activation (unstimulated T-cell population); PHA-activation (i.e., one day after PHA-stimulated); HIV infection (i.e., one day after HIV infection); as well as one and four days after xfR5-D+T NP or xfR5 mAb treatment (Figure 3).

**Figure 3.**
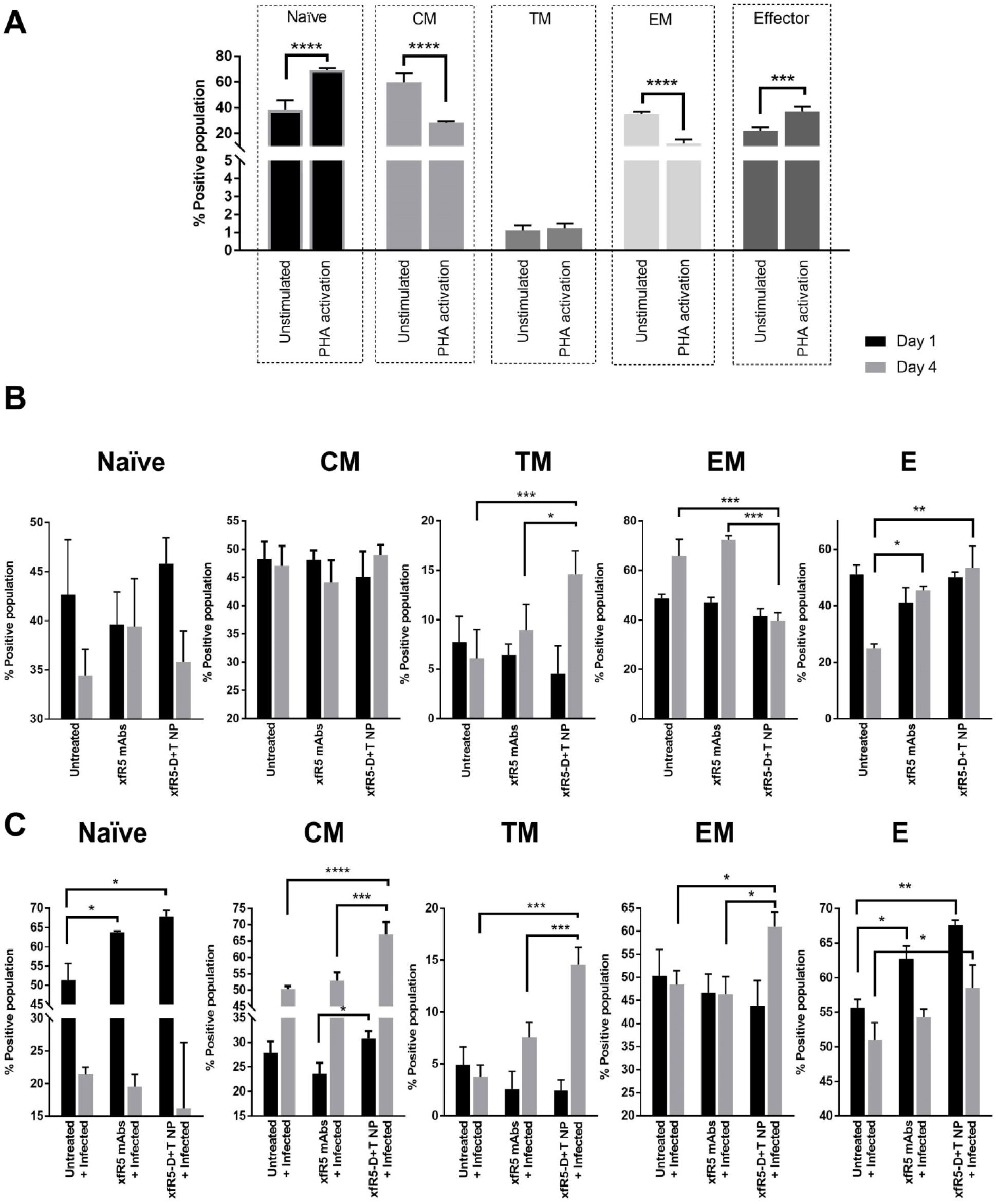
T-cell differentiation phenotype under untreated (A), PrEP (A), and treatment (B) condition. A) The untreated-nonactivated PBMCs were compared with PHA-activated PBMCs and immunophenotypic differentiation (after day 1) of naïve, CM, TM, EM, and E sub-population was evaluated. B) The immunophenotypic differentiation in untreated PBMCs compared to xfR5 mAbs and xfR5 D+T NPs were compared. The data presented as mean ± SEM of three independent experiments on three healthy donors (n=3). The significance was represented by the asterisk (*) symbol, where, ‘*’, ‘**’, ‘***’ and ‘****’ corresponding to *P* values <0.05. <0.01, <0.001 and <0.0001, respectively.

The unstimulated PBMCs evaluation was composed of 18%, 30.3%, 1.4%, 60.7% and 36.8% of naïve, CM, TM, EM, and E T-cell sub-populations, respectively (Figure 3A). As expected, the PHA stimulation (mock infection) significantly changed the immunophenotype of T-cell sub-populations stimulating the naïve and E sub-population (45.7% and 66.1%, respectively). Moreover, the EM population was significantly decreased (27.4%) during PHA stimulation. Emergence of activated effector cytotoxic T-lymphocytes (aCTLs, CD8+ CD69+) from minimal (0.66%) to 41.8% could be also be attributed to PHA-activation effect (Figure 4A).

**Figure 4.**
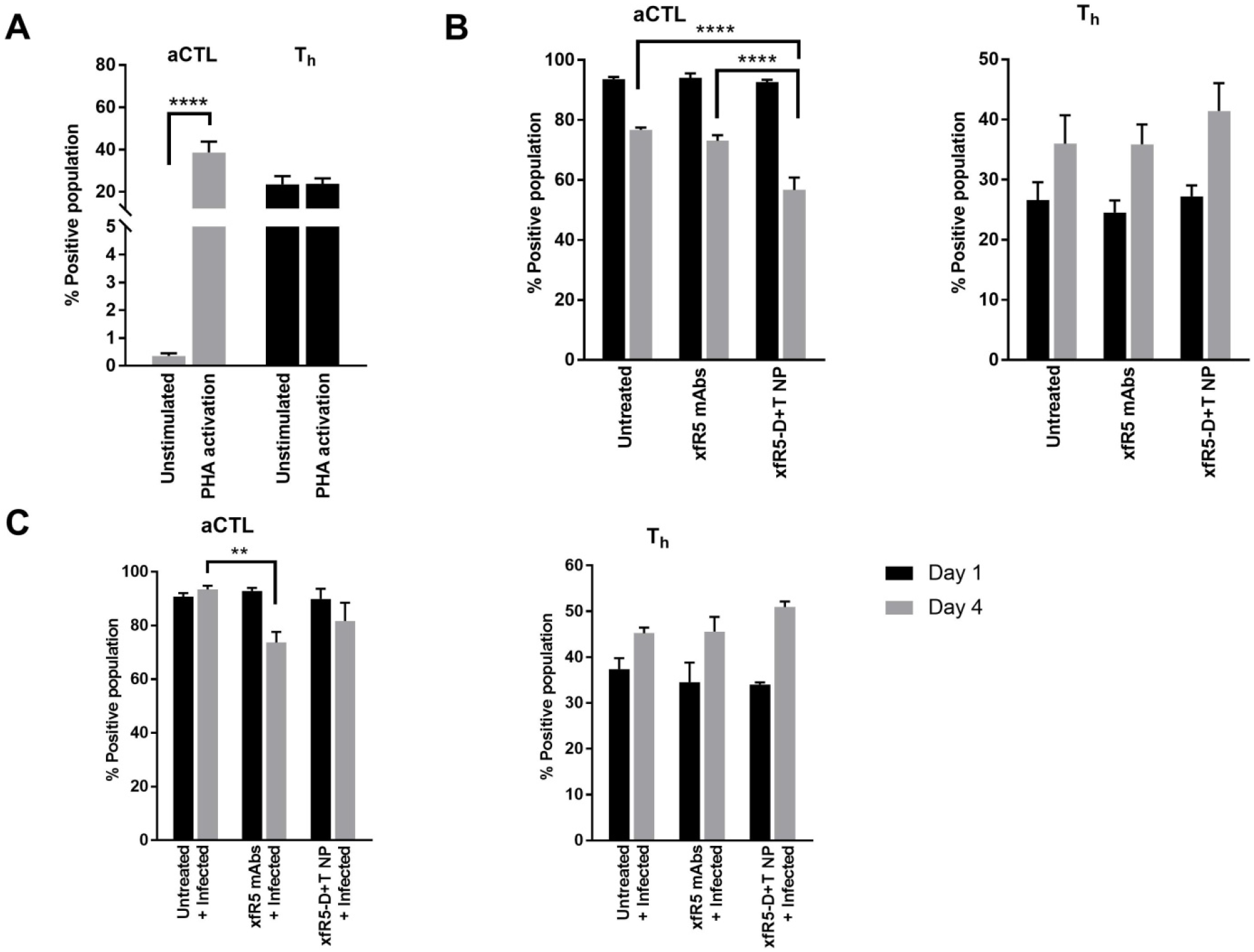
The effect on effector (E T-cell) phenotype upon treatment (with xfR5-D+T NP and xfR5 mAbs) compared to untreated conditions during PrEP and HIV-1 infection. (A) Comparative effect on aCTL (CD8+ E T--cell) and T_h_ (CD4+ E T-cell) phenotype under unstimulated and stimulated (PHA-activated). Under PrEP condition (B) aCTL (left graph) and T_h_ (right graph) population on day 1 (black bar) and day 4 (gray bar). (C) The treatment effect in the presence of HIV-1 infection, on aCTL (left graph) and T_h_ (right graph) on day 1, and day 4. Each data set is representing mean ± SEM of three independent experiments on PBMCs obtained from 3 healthy donors (n=3). The gating strategy for this study has been explained in Supplementary Figure 3. The data represented in the graph were mean ± SEM of three independent studies on three healthy donors (n=3). The significance represented as the asterisk (*) symbol, where, ‘*’, and ‘****’ corresponding to *P* values <0.05. and <0.0001, respectively.

The short-term PrEP (4 days) *ex-vivo* study, demonstrated xfR5-D+T NP significant protection against HIV infection (Table 4), which in part could be attributed to a protective immunophenotype (Figure 3B). The xfR5-D+T NP resulted in increased CM, TM, and E sub-populations. However, compared to PHA-activated (untreated), the xfR5-D+T NP demonstrated a significant boost in TM and E sub-populations (Figure 3B). The aCTLs and T_h_ population demonstrated a reciprocal effect (Figure 4B). The aCTL sub-population after initial spike (day 1) followed a decline; in contrast, T_h_ population showed a gradual increase in overtime (day 4) (Figure 4A, B).

The *ex-vivo* HIV-1 infection and short-term treatment (4 days) study were performed to predict the immunophenotypic differentiation pattern that the novel treatment could present (Figure 3C). The study showed the presence of HIV infection, mainly influencing the naïve and CM sub-population (Figure 3C). In the presence of HIV infection, the naïve sub-population after initial increased (day 1, post-infection, PI) reverts to basal levels (day 4, after HIV infection), and the CM population showed gradually increased population (day 4 PI). The xfR5-D+T NP treated population also showed enhanced naïve and CM sub-population during the initial active HIV infection stage (day 1 PI). In the long-term, however, the xfR5-D+T NP treated population demonstrated a reverse effect in the CM and TM sub-population. Prolong xfR5-D+T NP treatment (day 4, after treatment) of HIV-infected PBMCs (day 5 PI) displayed 2.4 times higher CM sub-population, as well as 6-times higher TM sub-population boost, compared to initial HIV infection. Moreover, xfR5-D+T NP treatment also significantly induced increases in EM sub-population. The xfR5-D+T NP treatment displayed significantly higher CM, EM, E, and T_h_ sub-population compared to xfR5 mAb and untreated controls. Therefore, these results demonstrate the multimeric interaction of xfR5-D+T NP could potentially improve the CM> TM > EM rate-limiting steps that would help in maintaining a high EM population during possible HIV challenge.

## Discussion

In this study, we have demonstrated a strategy to combine two different HIV treatments in a single nanoformulation with sustained release property. To achieve this we formulated CCR5 targeting ARV loaded nanoformulation with the potential to block HIV entrance and promote cellular immunity against HIV infection, specifically in memory CD4+ T-cells as they commit themselves to provide antiviral immunity as latent HIV-1 reservoir (34). Alongside memory CD4+ T-cells, the myeloid lineage, such as monocytes/macrophages, are believed to be other potential HIV-1 sanctuaries (35, 36). Our hypothesis is the novel CCR5 targeted ARV nanoformulation approach could combine different strategies to ensure dual protection against HIV to the CCR5+ cell types. First, CCR5 receptor docking on the CCR5+ cells will block HIV entrance to prevent HIV infection; and second, intracellular ARV released from endocytosed NP will providing protection against HIV-1 intracellularly to the naïve cell or latent HIV-infected cell population (Figure 1). The novel nanoformulation will also boost anti-HIV immunity to contribute to the possible “functional cure” strategies (37).

To target and block CCR5 receptors, the nanoformulation was surface decorated with high-affinity CCR5 mAb (xfR5 mAb) and D+T ARV were encapsulated within the nanoformulation. The w-o-w emulsion method produced uniform size xfR5-D+T NPs with a weak surface negative charge. The enhanced electron density (Figure 2B) and BCA protein quantification (Table 1) validated covalent binding of xfR5 mAbs were attached to the xfR5-D+T NPs (Figure 2C). Besides, the presence of amide bond (Figure 2C), and O–H stretching (3000-3500 cm-1) conferred xfR5 mAbs are surface decorated with an ample amount of free surface on NP with predominant PEG-PLGA composition. It is known that the onset of steric crowding compromises the binding avidity during multimeric interaction (38). Therefore, the spare xfR5 mAb coverage reduces the possibility of steric hindrance during multimeric xfR5 to CCR5-binding on the T-cells.

The observed enhanced binding affinity of xfR5-D+T NP (Table 2) could be attributed to the multi-valent interaction strengthening the binding affinity between the target biomolecules on the NP and the receptors on cell-surface (39). Previous theoretical calculation has illustrated that mAb conjugated to NP via flexible PEG spacers, could result in ten mAb/receptor complex formation per one NP interacting with respective receptor-expressing cells (39), although, one naïve xfR5 mAb in solution could only bind/block one CCR5 molecule per cell (32) (Supplementary figure 1). Therefore, based on the above theoretical assumption, the xfR5-D+T NP was expected to elucidate 10-times higher affinity than the naive xfR5 mAb. However, practically xfR5-D+T NP resulted in 7-times higher binding affinity compared to xfR5 mAb on TZM-bl cells.

The CCR5 co-receptor characteristically is expressed on CD4+ T-cells in peripheral blood (40). In addition to CD4+ T cells, the CCR5 chemokine receptor is also expressed on CTLs (CD8+) (41), dendritic cells (CD2+) (42) and monocytes/macrophages (CD68+) (43, 44), that also plays critical roles in HIV infection, propagation, and latency. Therefore, the binding affinity of xfR5-D+T NP on other CCR5+ expressing cell types are essential to estimate the success of this targeted nanoformulation strategy. Our study shows that the multimeric binding potency of xfR5-D+T NP evaluation on primary HIV-1 susceptible cells type, especially memory T_h_ (CD4+) and CTLs (CD8+) T-cell population showed ~5 and ~25 times higher binding affinity compared to naïve xfR5 mAb (Table 2). The enhanced CCR5 expression during infection or PHA-activation promotes migration of activated effector, and memory CTLs T-cells (45) Possibly enhanced CCR5+ expression on T-cells due to induction (by PHA-activation) of migrating phenotype possibly contributes to the substantial enhanced binding affinity of xfR5-D+T NP to CTLs. However, dendritic CD2+ populations and monocyte/macrophage CD68+ populations demonstrated slightly high binding affinity for xfR5-D+T NP compared to naïve xfR5 mAb. The reason behind the observed non-significant difference in binding affinity with dendritic CD2+ T-cells could be due to the low frequency of CCR5+ CD2+ the dendritic population in PBMCs (supplementary Figure 2) (46). Whereas, phagocytic CCR5+ CD68+ cells, i.e., monocytes/macrophages population (47), in addition to CCR5 receptor-specific binding, the non-specific phagocytotic uptake of xfR5-D+T NP and xfR5 potentially contributes to binding affinity evaluation.

The pronounced HIV protection efficacy of xfR5-D+T NP, as observed in the short-term PrEP study (Table 4), could be attributed to the enhanced binding affinity (Table 2) as well as sustained maintenance of intracellular ARV within xfR5-D+T NP treatment (Table 3). Further, the xfR5-D+T NP dual protection, i.e., multimeric xfR5 induced CCR5 blocking and intracellular ARV mediated HIV inhibition, boosted IC_50_ value by 526 (TZM-bl cells) and 12 (PBMCs) times compared to xfR5 mAb (Table 4). The combination or dual protection of xfR5-D+T NP demonstrates more rigorous protection compared to only blocking CCR5 receptors on T-cells by multimeric CCR5 T-cell blocking, i.e., 5.5 times higher; and ARV induced HIV protection by treating with D+T NP alone, i.e., 3.5 times compared to ARV by D+T NP alone. Further, it is well established (9, 10, 30, 31) and our study also reflects (Table 4), that reduced cytotoxic effect and the increased protection efficacy against HIV, widens the therapeutic index of xfR5-D+T NPs against HIV-1 virus. The enhanced high therapeutic index, therefore, advocates the potency of xfR5-D+T NPs as an advanced HIV therapeutic.

In general terms, the T-cell differentiation progresses in the following path: Naïve cells ⇒ CM ⇒ TM ⇒ EM ⇒ E. During initial HIV-1 infection, the immune system promotes naïve T-cell differentiation to antigen-specific memory and E sub-population (48). The immunophenotype evaluation (Figure 3) during the *ex-vivo* PrEP study demonstrated xfR5-D+T NP treatment enhanced naïve T-cell, memory (CM and TM), and E sub-population and the effect was sustained over the entire study period (Figure 3B). Therefore, the immunophenotype suggests that xfR5-D+T NP for PrEP application will keep the immune-system ready to promote fast clearance of the virions upon HIV challenge.

HIV-1 infection and ART demonstrated slightly different immunophenotype. Studies have shown that HIV-1 infection induces naïve T-cells proliferation, whereas ART causes depletion (49). A similar result was observed upon treating HIV+ PBMCs with xfR5-D+T NP and xfR5 mAb (Figure 3C). The *ex-vivo* immunophenotype of HIV-infected PBMCs demonstrated an increase in the naïve sub-population (47, 48). Whereas, xfR5-D+T NP treatment-induced HIV-1 suppression (Table 4) reverted the naïve sub-population to its basal level (similar to untreated+unfected+non-activated condition, Figure 3A). Moreover, the declined naïve T-cell population and increased CM sub-population could be due to differentiation induced population shifting from naïve CM sub-population.

The memory T-cell population plays a vital role not only in promoting anti-HIV immunity but also in AIDS prognosis since these cells govern the functions of CTLs and B-cells, which in turn regulates cellular and humoral immunity against HIV (50). Persistent HIV-1 viremia is known to drive the differentiation CM EM sub-population (51, 52), and the high EM sub-population may be responsible for the long-lasting and exhausted T-cell population in HIV+ patients (52). The ART treatment effectively partially restores high CM sub-population and declines EM sub-population over time to rebalance the homeostasis disrupted during HIV infection (52). This study shows as treatment progresses, xfR5-D+T NP treated population demonstrated significant increased CM sub-population (day 4, after treatment), as well as atypical to ART treatment, also significantly boosted TM and EM sub-population (Figure 3C). During active infection (Day 1 PI), targeted specific treatment (xfR5-D+T NP and naïve xfR5 mAb) maintained high E and aCTL sub-population, to counter HIV infection. Upon viral suppression (Table 4), E and aCTLs sub-population tend to decrease (Figure 4B). Studies have shown that despite persistently low antigenemia, high T_h_ and E sub-population is essential to control HIV replication in the presence or absence of ART to achieve “functional cure” (50, 53). However, long-term patients under ART-treatment results in the loss of effector functions. We observed HIV targeted treatment prolongs high E (Figure 3C) and T_h_ (Figure 4B) sub-population. Overall, the immunophenotypic study displayed the xfR5-D+T NP multivalent interaction results in enhanced and improves protective and defensive immunophenotype compared to naive xfR5 mAb treatment. Additionally, the multivalent phenomenon also contributes to maintaining high TM and EM sub-population. Therefore, the immunophenotypic differentiation study indicated that targeted cARV NP could also overcome the CM ⇒ TM ⇒ EM rate-limiting steps to promote high E and aCTL population maintenance during the HIV challenge.

In summary, we formulated a dual-action targeted nanoformulation, i.e., xfR5-D+T NP. The clinically relevant *ex-vivo* system (primary PBMCs) study showed xfR5-D+T NP induced CCR5 blocking and intracellular ARV drug release which provides two-levels of protection against fresh HIV-1 infection or in latently infected HIV+ cells. Further, xfR5-D+T NP treatment also promotes reprogramming of the immune repertoire function of memory T-cells and could help to reconstitute anti-HIV immunity. This novel targeted ARV nanoformulation promotes protective immune differentiation in T-cells. However, further investigations are essential to confirm the anti-HIV immune efficacy of xfR5-D+T NP, especially with HIV+ patients. However, based on this study, we conclude that xfR5-D+T NP combines the advantages of ART and augmentation of anti-HIV immunity reconstitution could be a promising multifactorial strategy to target the complex HIV latent reservoir to achieve HIV “functional cure.”

## Materials and methods

### Materials

Acid terminated PLGA (75:25, Mn = 4000–15000 Da), polyvinyl alcohol (PVA), dimethyl sulfoxide (DMSO), potassium dihydrogen phosphate (KH_2_PO_4_), dichloromethane (DCM), IL-2 and methanol were all purchased from Sigma-Aldrich (St. Louis, MO). Poly(lactide-co-glycolide)-block-poly(ethylene glycol)-succinimidyl ester (PLGA-PEG-NHS) and Methoxy Poly(ethylene glycol)-b-Poly(lactic-co-glycolic acid; PEG-PLGA (5,000:15,000 Da, 75:25 LA:GA) was purchased from Akina, Inc. (IN, USA). Pluronic F127, the stabilizer was obtained from D-BASF (Edinburgh, UK).

The ARV drugs, i.e., DTG (98% purity) was purchased from BioChemPartner co., Ltd., (China), whereas TAF (100% purity) was a generous gift from Gilead Sciences Inc. (CA, USA), under MTA agreement. Internal standards i.e. tenofovir-d6 (TFV-d6), TAF-d5, TFV-dp-d6 and dolutegravir-d4 (DTG-d4) were purchased from Toronto Research Chemicals Inc. (ON, Canada).

Roswell Park Memorial Institute (RPMI) 1640 with L-glutamine medium, Dulbecco’s Modified Eagle Medium (DMEM) high glucose medium (HiDMEM) and 100 × antibiotic-antimycotic (AA) were purchased from ThermoFisher Scientific (OK, USA), whereas fetal bovine serum (FBS) was from VWR International (PA, USA). All the chemicals were used as received.

### Primary cells and cell lines

The TZM-bl cell line obtained from the National Institutes of Health (NIH) acquired immunodeficiency syndrome (AIDS) reagent program is a JC53-bl (clone 13)/HeLa cell line that are phenotypic similar to HIV-1 infecting cell type (stably overexpresses CD4 and CCR5 receptor). TZM-bl cells were maintained in HiDMEM supplemented with 10% FBS and 1× AA, as standard protocol (9, 10). Whereas, peripheral blood mononuclear cells (PBMCs) were purchased from AllCells^®^ (Alameda, CA, USA) and maintained in RPMI 1640 supplemented with 10% FBS, 1×AA and 50 U/ml IL-2. The XF-CCR5 28/27 43E2AA hybridoma (xfR5 mAb producing cell line), was maintained in RPMI supplemented with 10% FBS and 1×AA (54).

### HIV strain

HIV-1_ada_ virus obtained from the NIH AIDS research program and further propagated by the following published standardized method (55). The TCID_50_ evaluated by standardized p24 ELISA assay using ZeptoMetrix^®^ HIV Type 1 p24 Antigen ELISA kit (NY, USA) on PBMCs received from healthy donors and following the manufacturer’s protocol.

### Production, purification, and characterization of xfR5 mAb

The modified high affinity xfR5 mAb were produced and isolated from a Hybridoma cell line, i.e., XF27/28/CCR5 43E2-AA (PTA-4054; ATCC repository) (54), by following published method with modifications as described below (56). Briefly, XF-CCR5 28/27 43E2AA hybridoma cells were seeded (at 10^6^/mL concentration) and maintained in antibody production inducing media, i.e., RPMI medium with 1×AA, for several days until 50% cells were found to be compromised (cell death). The supernatant with soluble xfR5 mAb was harvested by pelleting out dead cells and debris. The soluble xfR5 mAb from the supernatant was isolated using HiTrap™ Protein-A HP prepacked column (GE Healthcare; NJ, USA) following standard manufacturer’s protocol. The purity and concentration of the xfR5 mAbs were determined respectively by the SDS-PAGE method and BCA assay using Pierce™ BCA Protein Assay Kit, following manufacturers’ protocol. The xfR5 binding was evaluated based on the standard curve (linear regression analysis) from a known concentration of IgG4 isotype control mAb, as xfR5 is recombinant human CCR5 mAb with IgG4 isotype backbone (54).

### D+T NPs formulation, xfR5 mAbs conjugation, and characterization

The targeted nanoformulation was fabricated by following multiple steps. First, NHS functionalized D+T loaded nanoformulation was obtained by following the modified oil-in-water emulsions phase inversion method (9, 10). Briefly, in the DCM organic phase PLGA, PLGA-PEG-NHS, PEG-PLGA, PF127 along with TAF and DTG have dissolved at 1:1:2:2:4:4 ratios, TAF (at comparative ratio 2) in PBS were added dropwise under constant stirring condition. The water-in-oil (w-o) emulsion was sonicated as described below and then added dropwise to the three times higher volume of 1 % PVA solution (aqueous phase) under high-speed stirring conditions. The above w-o-w emulsion was immediately probe-sonicated for 5 mins on ice (setting: 90% Amplitude; pulse 0.9 cycle/bursts) with the help of UP100H ultrasonic processor (Hielscher Inc. Mount Holly, NJ, USA). The organic phase from the o-w emulsion was completely evaporated overnight (O/N). The NHS functionalized D+T NPs were desiccated by lyophilization using Millrock LD85 lyophilizer (Kingston, NY, USA) to eliminate the aqueous phase. The complete formulation method was carried out under the hood to maintain sterility during fabrication.

The NHS functionalized D+T NPs, xfR5 mAb was conjugated using its amine group to amine-reactive NHS esters of D+T NPs (57). Briefly, NHS ester-D+T NPs were disassociated in PBS (pH 7.4) as the NHS esters (4-5 h half-life of at pH 7.4). The xfR5 mAb maintained in PBS to have protonated amine groups of xfR5 mAb. The xfR5 mAb were added to NHS ester-D+T NPs at 1:10 weight ratio under constant stirring at room temperature (RT), and the reaction proceeded for 2 hours at RT. Immediately, after the reaction, the unbound xfR5 mAb was washed off by dialysis using Float-A-Lyzer^™^ G2 Dialysis Devices (Thermo Fisher Scientific; NH, USA) in cold PBS supplemented with 10mM hydroxylamine to quench any non-reacted NHS groups present on the xfR5 mAb conjugated D+T NPs (xfR5-D+T NPs) surface. Followed by three consecutive cold PBS buffer-exchanges, xfR5-D+T NPs were collected and stored at 4°C. The concentration of xfR5 mAb conjugated on the xfR5-D+T NPs surface was evaluated by the BCA assay. To avoid background issues, PBS, as well as D+T NPs, were run parallel during BCA assay, and the obtained values were considered background.

The physicochemical properties of the D+T NPs were evaluated based on dynamic light scattering (DLS), fourier transform infrared (FT-IR) spectroscopy, and scanning electron microscopy. The size, surface charge, and polydispersity index (PDI) of the D+T NPs were determined by a ZetaPlus Zeta Potential Analyzer instrument (Brookhaven Instruments Corporation; NY, USA) following standardized methodology (9, 10). The DLS analysis determined the D+T NPs size and polydispersity index (PDI), i.e., size homogeneity and size-distribution pattern. The zeta potential analysis identified the surface charge density on the D+T NPs. The D+T NP’s surface NHS functionalization and xfR5 mAb binding via amide bond were evaluated based on FT-IR spectroscopic analysis following a previously published method (58). Briefly, the spectra of each sample in powder form were collected in the range 600–4000 cm^-1^ using 25 scans at a resolution of 4 cm^-1^ with % transmittance intensity mode and Happ-Genzel function apodization under IRPrestige-21 Fourier transform infrared spectrometer (FT-IR) instrument (Shimadzu; MD, USA) and by using LabSolutions IR software (Shimadzu; MD, USA) the data were analyzed. By scanning electron microscopy, the morphology and shape of the D+T NPs were evaluated (59). Briefly, D+T NPs were deposited on Whatman^®^ Nuclepore Track-Etch Membrane (~50 nm pore size) and air-dried for one day at RT under a chemical hood. The air-dried NPs membrane was sputter-coated with a thin layer (~3-5 nm thick) of chromium and imaged under a Hitachi S-4700 field-emission SEM (New York, NY, USA).

The % drug entrapment efficiency (%EE) of DTG and TAF in D+T NPs and xfR5-D+T NPs were evaluated by high-performance liquid chromatography (HPLC) instrument by following published methodology (30, 31, 60). Briefly, 1 mg of D+T NPs dissociated in 50 μL DMSO and mobile phase (25mM KH_2_PO_4_ 45%: ACN 55%) added to get 10% DMSO final concentration in the injection volume (20 μL). For the standard curve evaluation, the same procedure was followed to prepare the D+T standard solutions (with each drug concentration from 0.5 to 0.0019 mg/mL). The chromatography separation was performed under a HPLC instrument (Shimadzu Scientific Instruments; MD, USA) equipped with SIL-20AC auto-sampler, LC-20AB pumps, and SPD-20A UV/Visible detector, using Phenomenex^®^ C-18 (150×4.6 mm, particle size 5 μm) column (Torrance, CA, USA), under isocratic elution process with 0.5 mL/min mobile phase flow rate, temperature: 25°C; and detection at 260 nm (retention time of 4 mins for TAF and 6.3 mins for DTG). The quantification of the drug was determined by evaluating the peak area under the curve (AUC) analysis at their respective retention time. The amount of TAF and DTG loaded in the D+T NPs was analyzed based on the standard curve construction (linear correlation, r^2^≥0.99) respective from TAF, and DTG standard concentration ranges from 0.5 mg/mL to 0.0019 mg/mL. The HPLC instrument illustrated inter-day and intra-day variability of <10%. The % encapsulation efficiency (%EE) of each drug in the D+T NPs batch was estimated by equation 1, respectively. The data presented as mean± standard error of the mean (SEM) of three D+T NPs batches (n=3).

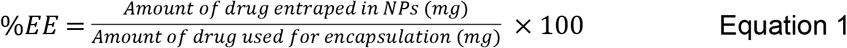

### Antibody binding and binding affinity evaluation

To establish and estimate the binding affinity of isolated xfR5 mAb and xfR5-D+T NPs in comparison to wild-type CCR5 mAb, Cy3 conjugated to xfR5 mAb by Cy3 NHS Ester Mono-Reactive CyDye (GE Healthcare; PA, USA), following manufacturer’s protocol. The Cy3 dye to xfR5 mAb binding ratio was evaluated based on regression analysis of respective standards (i.e., 0.5 to 0.00625 mg/mL) data obtained from UV/vis spectroscopy and BCA assay. The 3:1 dye to xfR5 mAb ratio Cy3 conjugated xfR5 mAb (Cy3-xfR5 mAb) batches considered for further studies. To study binding affinity by flow-cytometry, Cy3-xfR5 mAb conjugated D+T NPs (Cy3-xfR5-D+T NPs) formulation were fabricated and characterized, similarly as described for xfR5-D+T NPs. By using the standardized formulation, three independent batches were obtained and evaluated for further studies.

For binding affinity of Cy3-xfR5 mAb and Cy3-xfR5-D+T NPs, TZM-bl cells (10^5^/well) and phytohemagglutinin (PHA, at 5 μg/mL) activated PBMCs (10^6^ cells/well) were treated with wild-type Cy3-CCR5 mAb (rabbit anti-human mAb; Bioss Inc.; MA, USA), Cy3-xfR5 mAb and Cy3-xfR5-D+T NPs at different concentrations (20, 10, 1, 0.1, 0.1 μg/mL of xfR5 concentration) O/N at 37°C and 5% CO_2_ atmosphere. Reportedly, PHA is known to induce reactivation of latent HIV-infected primary T-cells [42] consistently, therefore, to achieve latent HIV-infected primary T-cells phenotype (promote CD2+ T-cells), PHA-activated PBMCs were used. The treatment was washed-off by washing thrice with 1% BSA in PBS (PBA) solution by centrifugation (220 × g at 4°C). As HIV-1 primarily infects CD4 T-cells, therefore, evaluate binding affinity of xfR5 mAb and xfR5-D+T NPs compared to wild type anti-CCR5 mAbs, all the above-treated cells were incubated with anti-CD4 AlexaFluor700 mAb (Table 5) for 20mins at RT (at 1:100 dilution) and washed thrice with PBA.

**Table 5.** Detailed information about different T lymphocyte phenotypic markers used for the immunophenotypic study.

| **T-lymphocyte target** | **Marker** | **Fluorescence dye conjugated** | **Monoclonal antibody type** | **Company** |
| --- | --- | --- | --- | --- |
| T lymphocytes | CD3 | Alexa Flour 594 | Mouse anti-human | BioLegend |
| helper T-cells | CD4 | PerCP-Cy5.5 | Mouse anti-human | TONBO biosciences |
|  |  | Alexa Flour 700 | Mouse anti-human | BioLegend |
| Cytotoxic T-cells | CD8 | PE/Cy7 | Mouse anti-human | BioLegend |
| Monocytes | CD68 | APC | Mouse anti-human | Invitrogen |
| HIV target on T-cell | xfR5 | Cy3 | Modified anti-human | ATCC |
| Transition T-cells | CCR7 | CCR7-APC eFlour 780 | Mouse anti-human | Invitrogen |
| Memory T-cells | CD45RO | Pacific Blue | Mouse anti-human | BioLegend |
| Activated T-cells | CD69 | Alexa Flour 488 | Mouse anti-human | BioLegend |
| HIV maker on T-cells | CCR5 | APC | Mouse anti-human | BioLegend |
| Intermediate memory T-cells | CD27 | Alexa Flour 700 | Mouse anti-human | BioLegend |
| DC or latently infected T-cells | CD2 | Pacific Blue | Mouse anti-human | BioLegend |

Similarly, to evaluate binding affinity with the latent population (CD2+ T-cells) and monocytes (CD68+ T-cells), PBMCs were treated as mentioned above. The above-treated cells incubated for 20 mins with anti-CD68 APC mAb and anti-CD2 Pacific Blue (Table 5). The above marker antibody bound treated cells were fixed for 20 min with 4% PFA at 4°C and washed again twice with PBA. The binding of Cy3-xfR5 mAb and Cy3-xfR5-D+T NPs to respective T-cell type was detected and evaluated respectively by the BD LSRII flow cytometer instrument (BD Biosciences; San Jose, CA, USA) and Flowjo software v10 (BD, Franklin Lakes, NJ, USA). The supplementary figure 1 details the complete gating strategies. Each study mentioned above was performed on three healthy independent donors PBMCs. The binding affinity was calculated based on Michaelis-Menten’s non-linear fitting analysis of mean ± SEM (standard errors of means).

### Immunophenotype study

Immunophenotype variation upon xfR5-D+T NPs treatment compared to xfR5 mAb in uninfected (mimicking PrEP condition) and HIV-1_ADA_-infected PHA-activated PBMCs was evaluated by flow cytometry. Briefly, PBMCs (10^5^ cells/well) were treated respectively with xfR5 mAb and xfR5-D+T NPs (at 20 μg/mL of xfR5 concentration) for 96 h at 37°C and 5% CO_2_ atmosphere. As the control and to compare activated PBMCs immunophenotype, PHA-activated PBMCs (10^5^ cells/well) were maintained alongside for 96 h (61). Treated cells were washed thrice with PBA solution by centrifugation (220×g at 4°C) and incubated with mAb against T-lymphocytes (CD3), helper T-cells (CD4), cytotoxic T-cells (CD8), memory T-cells (CD45RO), transition T-cells (CCR7), activated T-cells (CD69), intermediate memory T-cells (CD27), and HIV latently infected T-cells (CD2) markers (Table 5), for 20 mins at RT (at 1:100 dilution). The marker-bound treated cells were washed with PBA, fixed for 20 min with 4% PFA at 4°C, and rewashed thrice with PBA. The immunophenotype of the markers bound treated cells were evaluated by flow cytometry. Three independent studies have been performed on three healthy donor’s PBMCs. The data presented as mean ± SEM obtained from three independent donors.

### Intracellular kinetics study

The intracellular uptake and retention kinetics of D+T NPs and D+T solution were evaluated by LC-MS/MS analysis following a standardized method (9, 30, 62). Briefly, TZM-bl cells (10^4^ cells/well) seeded in the 24-well plate with the complete HiDMEM medium. Following O/N cell adherence, respective cells group were treated with D+T NPs and D+T solution at 10 μg/mL concentration of each drug, i.e., DTG and TAF. For uptake kinetics study, at respective time-points (i.e., 1, 6, 18, and 24 h), the treated cells were then washed twice with warm PBS and detached by Trypsin-EDTA (25%; Thermo Scientific, OK, USA), washed with twice with PBS. One set of untreated detached cells were counted at each time-point to determine cell count at respective time-point. The cells were air-dried under a biosafety cabinet. The air-dried samples were then lysed with 70% methanol and stored at −80°C until analysis. Whereas, for the drug-retention kinetics study, the adhered TZM-bl cells were treated with xfR5-D+T NP and D+T solution, respectively, for 24 h and washed thrice with warm 1×PBS. The washed treated cells were in fresh complete HiDMEM medium until respective time-points (i.e., 1, 6, 24, and 72 h after wash, that corresponds to 25, 30, 48, and 96 h, time-point respectively after treatment). At respective time-point, the cells were rewashed with PBS, detached, lysed, and stored following the same method as explained above. The samples were analyzed using the LC-MS/MS method described in the section below.

For the intracellular DTG, TAF, TFV, and TFV-dp drug-kinetics evaluation by LC-MS/MS instrument, the respective cell lysates were centrifuged (14000 rpm for 5 mins at 4°C) and the supernatant was collected. To an aliquot of 100 μL supernatant, 300 μL of internal standard spiking solution (10 ng/mL each of DTG-d4, TAF-d5, TFV-d6, and 100 n/mL of TFV-dp-d6 in ACN) was added, and vortexed. The samples were then dried at 45°C under the stream of nitrogen and reconstituted with 100 μL 50% acetonitrile. The drug and metabolites were quantified from the same sample using LC-MS/MS instrument.

For TAF, TFV, and DTG estimation, the similar conditions that were previously published by our group were used with minor modification (62). One μL of the processed sample was injected on to LC-MS/MS operated in positive mode. The chromatographic separation was carried-out using the Restek Pinnacle DB Biph column (2.1 mm × 50 mm, 5 μm) with 0.5% formic acid in water and 0.1% formic acid in ACN (48:52 v/v) mobile phase. The calibration range for all the analytes was 0.01 to 50 ng.

For the quantification of TFV-dp, Phenomenex Kinetex C18 (75×4.6 mm, 2.6μm) column was used with an isocratic mobile phase (10mM ammonium acetate pH 10.5: ACN (70:30) at a flow rate of 0.25 mL/min. The dynamic calibration range was from 0.01 to 100 ng. The LC-MS/MS system consisting of an Exion HPLC system (Applied Biosystems, CA, USA) coupled with AB Sciex 5500 Q Trap with an electrospray ionization (ESI) source (Applied Biosystems, CA, USA) was used in positive ionization mode. The retention time of TFV-dp was 2.1 min, and the runtime for each sample was 3.5 min. The average inter-day and intra-day variability were < 15%, which corresponds to the FDA bioanalytical guidelines (63).

### *In vitro* cytotoxicity Study

The comparative *in vitro* cytotoxicity of D+T NP vs. D+T solution was evaluated on the TZM-bl cell line using CellTiter-Glo^®^ luminescent assay method, as described previously (64). Briefly, the TZM-bl cells (10^4^ cells/well) in complete HiDMEM medium and PBMCs (10^5^ cells/well) in complete RPMI (Thermo Scientific; OK, USA) supplemented with 10% FBS, 1 × AA and 50 U/ml IL-2 (Sigma-Aldrich; MO, USA), were treated in triplicate respectively with D+T NP or D+T solution, at different concentrations (20, 10, 1, 0.1, 0.01 μg/mL each drug concentration) for 96 h. Similarly, the 5% DMSO treated cells and 1×PBS (treatment equal volume) treated cells were the positive and negative control, respectively. The cytotoxicity was evaluated by the CellTiter-Glo^®^ luminescent-cell viability assay kit (Promega; WI, USA) following manufacturer protocol. The luminescence intensity read under the Synergy II multi-mode reader with Gen5TM software (BioTek; VT, USA) and the percentage cytotoxicity (% cytotoxicity) values obtained by subtracting the % normalized viability (against the untreated negative control group) from 100. The experiment was carried out on three independent batches of D+T NPs and D+T solution. The result represents the mean ± SEM of three independent batches studies. The untreated control cells were considered 100% viable. The % cytotoxicity was evaluated by following equation 2:

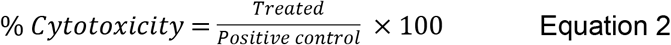

### *In vitro* protection study

The comparative *in vitro* prophylaxis (PrEP), i.e., protection study between D+T NP vs. D+T solution against HIV-1_NL4-3_, was performed on TZM-bl cells (an HIV-1 infection reporter cell line), and peripheral blood mononuclear cells (PBMCs) was evaluated by following standardized method (9, 64). Briefly, TZM-bl cells (10^4^ cells/well) and PBMCs (10^5^ cells/well) were seeded in 96-well plate and were treated with different concentrations of D+T (20, 10, 1, 0.1, 0.01 μg/mL each drug concentration) either as D+T NP or as D+T solution. Whereas untreated/uninfected cells and untreated/infected cells were considered negative and positive controls, respectively. After 24 h of treatment, the TZM-bl cells were infected with HIV-1_NL4-3_ virus (multiplicity of infection, MOI: 1) for 8 h, whereas PBMCs got HIV-1_ADA_ virus (MOI: 0.1) infection for 16 h. At respective time-point, the inoculated and control cells were washed with warm PBS (thrice). The TZM-bl cells and PBMCs were then maintained respectively in fresh complete HiDMEM medium and complete RPMI for 96 h. The HIV-1 infectivity was evaluated by the luminescence intensity following Steady-Glo^®^ luciferase assay (Promega; WI, USA), following company specified methodology. The luminescence intensity based on relative luminescence units (RLU) due to HIV-1 infection, was read on the Synergy HT Multi-Mode Microplate Reader (BioTeck; Vt, USA). The % HIV-1 infection was calculated by following equation 3:

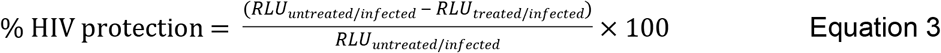

For TZM-bl cells, data collected from three independent experiments performed with three different batches of D+T NP and D+T solution (each performed in duplicate). For PBMCs, the data obtained after treating three different healthy donor’s PBMCs (each performed in triplicate) at independent time. Finally, the selectivity index (SI), was evaluated by following equation 4:

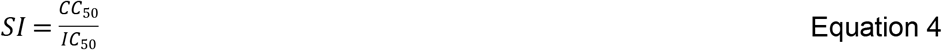

Where, ‘CC_50_’ (cytotoxic concentration at 50%) and ‘IC_50_’ (50% inhibitory concentration), was evaluated from the above described *in vitro* cytotoxicity and protection study.

### Statistical Analysis

All study results presented are expressed as mean ± SEM of the obtained data from at least three independent experiments. The CC50 value was determined by non-regression curve fitting based on log (DTG or TAF concentration) vs. normalized luminescent (three-parameter logistic fits) of cytotoxicity response curves. Whereas, the IC_50_ value was evaluated by fitting the non-regression inhibitory curve of log [DTG] vs. normalized TZM-bl luminescence (three-parameter logistic fits) luminance values. Analysis of variance (ANOVA) method was used to determine significant differences between treated (D+T NP and D+T solution) vs. control groups at p-value ≤ 0.05. All the statistical analysis presented was determined by GraphPad Prism 7 software (La Jolla, CA, USA).

## Acknowledgment and Support

These results presented were funded by the State of Nebraska Cigarette Tax (LB692) grant, and NIAID R01AI117740-01, 2015 to C.J.D. Authors are thankful to Gilead Sciences, Inc. for providing TAF drug under MTA. A special thanks to the UNMC Flow Cytometry Research Facility (UNMC FCRF). The UNMC FCRF is administrated through the Office of the Vice-Chancellor for Research and supported by state funds of the Nebraska Research Initiative (NRI) and the Fred and Pamela Buffett Cancer Center’s National Cancer Institute Cancer Support Grant.

